# A theoretically driven calculation for language dominance and degree of multilingualism

**DOI:** 10.1101/2025.06.13.659417

**Authors:** Xuanyi Jessica Chen, Esti Blanco-Elorrieta

## Abstract

Bilingualism research has long been challenged by a lack of a unified approach to quantifying language dominance and degree of multilingualism. While numerous questionnaires (e.g., LHQ, BLP, LEAP-Q, and LUQ) provide valuable data on language background variables, they lack a standardized formula to compute key measures from it. We introduce two formulas that synthesize critical linguistic variables to efficiently calculate language dominance and a multilingualism score that ranges from perfect monolingualism to native-like proficiency in multiple languages. Validation across two large datasets shows our dominance measure closely aligns with more complex PCA methods while being simpler and more efficient.

## 1 Introduction

One of the biggest challenges in bilingualism research is accurately determining relative language dominance from language background data. While standardized questionnaires exist for collecting such data, there is no widely accepted formula for calculating dominance. These questionnaires differ in focus and detail, with commonly used examples including the Language History Questionnaire (LHQ) (Li et al., 2014), Bilingual Language Profile (BLP) (Birdsong et al., 2012), Language Experience and Proficiency Questionnaire (LEAP-Q) (Marian et al., 2007), and Language Use Questionnaire (LUQ) (Kastenbaum et al., 2019; Marte et al., 2025). They typically assess factors such as language exposure, proficiency, age of acquisition, and frequency of use to estimate bilinguals’ relative proficiency (e.g., Blumenfeld & Marian, 2013; Grant & Dennis, 2017; Mercier et al., 2014; Titone et al., 2011). However, despite efforts to gather increasingly comprehensive linguistic background data, the challenge of quantifying language dominance and the degree of multilingualism from this data remains unresolved.

Some of the primary methods used to compute language dominance in the literature include A) Self-reported dominance scores (e.g., Birdsong et al., 2012; Marian et al., 2007), which ask bilinguals to rate their dominance on a Likert scale, relying on subjective judgments (Amengual, 2016; Gollan et al., 2007; Lemhöfer et al., 2004; Paap & Greenberg, 2013). B) specific performance metrics, where researchers assess dominance using vocabulary size, lexical retrieval, morphological and syntax competence, or verbal fluency measures (e.g., Bonvin et al., 2021; Gollan et al., 2012; Paradis & Nicoladis, 2007; Pienemann et al., 2011). C) Principal Component Analyses (PCA) to reduce all language background variables to a single dimension (e.g., Kastenbaum et al., 2019; Chen et al., submitted) or D) Average scores, which vary on whether they exclusively include language background variables or whether they combine language experience measures (e.g., proficiency) with some objective measure. For example, Robinson and Blumenfeld (2019) created a composite dominance score by combining two subjective measures—self-reported language abilities (speaking, understanding, listening, and reading) and exposure to Spanish and English—with an objective proficiency score from the MINT (Gollan et al., 2012; see also Gálvez-McDonough et al., 2024).

While these approaches have been important steppingstones, they (i) assign equal weight to every background measure despite the likelihood that their contributions to language dominance vary, and (ii) do not incorporate established insights about bilingual language acquisition into these calculations. Importantly, the heterogeneity in these approaches highlights a critical issue in the field: despite many bilingualism studies collecting extensive language background data, there is no clear, standardized method to synthesize this information into a single measure of language dominance or degree of multilingualism, which makes it difficult to apply the collected data in systematic ways during analysis and interpretation. Further, without a unified formula, the field risks obtaining inconsistent or non-comparable results, undermining the generalizability of results across bilingualism studies.

Thus, a more precise and theoretically grounded approach to measuring language dominance is needed. The formulas proposed in this paper synthesize key linguistic variables while eliminating redundancy in the calculations, providing a time-efficient and replicable assessment of bilingual experience.

## 2 Methods

### 2.1 Language Dominance measure

The goal of computing a language dominance measure is to obtain a continuous score ranging from −1 (fully dominant in one language) to 1 (fully dominant in the other), with 0 indicating a perfect balance between both languages. Here, we applied the following approach to obtain this value in a streamlined, theoretically informed manner.

#### Step 1: Deriving Normalized Proficiency Scores

For each participant and language, we averaged self-rated scores for listening, speaking, writing, and reading into a composite proficiency score, then normalized it between 0 and 1 relative to each participant’s highest rating to account for individual biases in self-reporting and ensure that a participant’s strongest language received a perfect proficiency score.

Here, we rely on self-rated proficiency over standardized test measures. This choice does not affect the nature of the formula or dominance calculation; it is driven by an accuracy-cost trade-off given that self-rated proficiency is highly correlated with objective measures such as the MINT (Marian et al., 2007; Gollan et al., 2012; Luk & Bialystok, 2013) and does not appear to provide additional explanatory power.

We tested this directly analyzing data from a sample of n = 38 (Blanco-Elorrieta et al., 2020) which contained both self-rated proficiency and standardized scores in speaking, hearing, reading, writing using the Woodcock–Muñoz English Language Survey (Woodcock et al., 2005). Variance analyses showed that self-rated ability accounted for over 97% of variance in standardized scores across all language abilities (Speaking = 97.8%; Listening = 98.3%; Reading = 98%; Writing = 97.9%), confirming that self-assessments effectively capture language proficiency.

Thus, given that standardized testing imposes practical costs including increased time, resources, and participant burden/stress; while offering little additional explanatory value beyond self-rated proficiency, we advocate for the most parsimonious and least labor-intensive approach.

#### Step 2: Modeling AoA Effects on Ultimate Proficiency Potential

To capture how ultimate proficiency attainment is modulated as a function of age of first exposure, we transformed the averaged AoA across the four language components using the following logistic function:

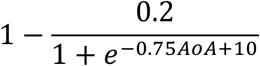

The parameters for this function have been derived from Hartshorne et al. (2018) and they reflect that individuals who acquire a language before age 10 can reach full native-like proficiency and thus retain a scaling value of 1. Beyond this age, the potential for native-like attainment gradually decreases, tapering off until around age 18, after which proficiency is capped at 80% of native-like ability. Vanhove (2013) noted that accurately estimating how ultimate proficiency changes with age of first exposure requires samples in the thousands, whereas studies typically relied on much smaller datasets (∼50 to 250 participants). To ensure robustness, we based model parameters on Hartshorne et al. (2018), the largest study to date with hundreds of thousands of bilinguals. Importantly, this function does not model the critical period for language acquisition thought to peak around age 7 (Johnson & Newport, 1989). Instead, it models how the age of first exposure will determine ultimate proficiency.

Having modeled the effect of Age of Acquisition on attained proficiency, we then multiply this AoA factor with the scaled ability from Step 1 to obtain a composite proficiency score for each language.

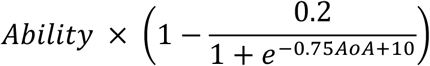

#### Step 3: Calculating Language Dominance

Finally, language dominance is defined as the difference in scaled proficiency between the pair of languages of interest (denoted as L1 and L2 in the formula):

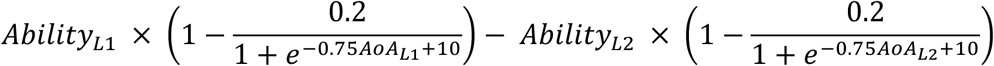

This value ranges from −1 to 1, where a value closer to 1 or −1 indicates a strong dominance in one language over the other, whereas values closer to zero indicate balanced bilingualism. For multiple examples of Language Profile simulations and corresponding Language Dominance calculations see Figure 1.

**Figure 1.**
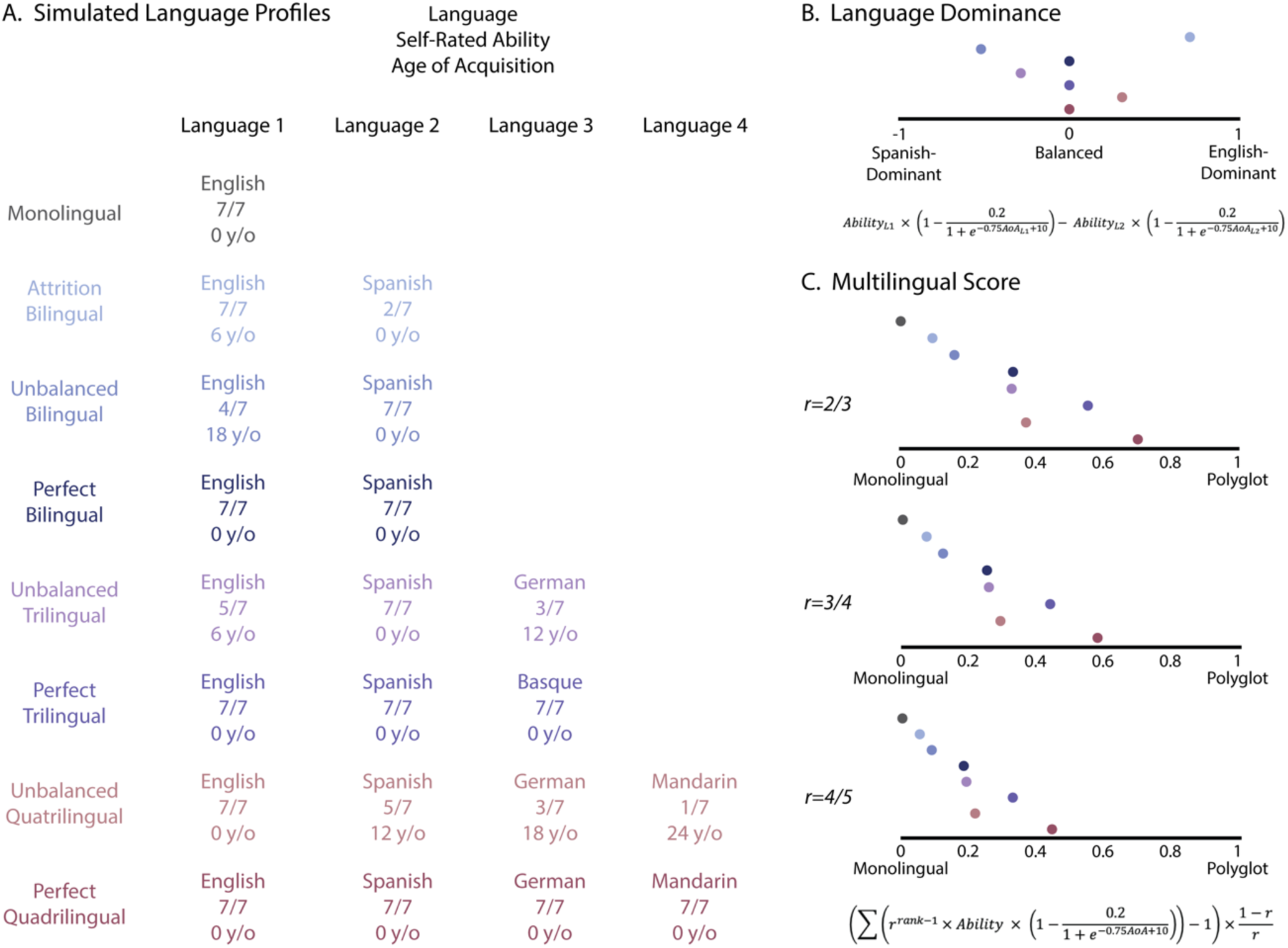
Language Dominance and Multilingual Score in simulated language background profiles. A. Simulated language profiles with the languages they speak, self-rated ability, and age of acquisition. B. Language dominance measure for each of the simulated profiles. C. Multilingual score with different common ratios that scales the weight of each additional language.

### 2.2 Multilingualism Score

Alongside a dominance formula, the field’s shift toward viewing bilingualism as a spectrum calls for a continuous measure that precisely quantifies degree of bilingualism. We propose an efficient multilingual scale that takes each participant’s ability rating and AoA in each language and outputs a value from zero to one, where zero corresponds to a perfect monolingual, and one to a hypothetical perfect polyglot. Steps 1 and 2 are as in the Language Dominance Calculation.

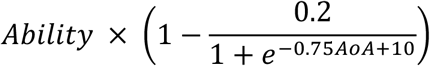

#### Step 3: Calculate each language’s weight

Each language’s weighting reflects the differential contribution of adding an additional language to one’s language experience, where, intuitively, the distance between a perfect monolingual and a perfect bilingual should be more pronounced than the distance between a perfect quadrilingual and a perfect quintilingual.

To compute this weighing, we first ordered each participant’s languages from their most proficient language (L1) to their least proficient language (Ln). If two languages had identical proficiency scores, their ranking was considered indistinguishable, and their order was randomized. Once the languages had been ranked, we had to determine the function by which the weight of each additional language would decrease. However, unlike the effect of AoA, the precise magnitude of this effect is empirically undefined. Here, we decided to use a geometric sequence in the form of *r^n-1^* where the initial value is 1 and the consecutive values are always scaled by the same ratio (*r*, the common ratio). The ratio should be between 0 and 1 so that the weights are always positive while decreasing in magnitude. This is theoretically motivated to ensure that each additional language carries progressively less weight than the previous one by the same proportion.

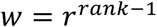

#### Step 4: Compute the final multilingualism score

To compute the final multilingual measure, we compute a weighted sum of language proficiencies. Then, we scale the resulting scores from 0 to 1. The unscaled weighted sum has a minimum value of 1 for monolingual speakers and approaches 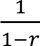 for a hypothetical polyglot that speaks infinite languages perfectly. This means that the total range of unscaled values is 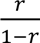. To scale the measure appropriately, we first subtract 1 to set the monolingual score to 0. Then, we multiply it by the inverse of the total possible range, 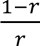 ensuring the values are normalized within a 0 to 1 scale. The resulting weight function can be expressed as below.

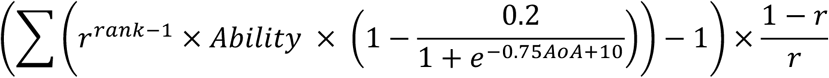

Here we tested three values for the common ratio: 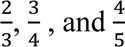, corresponding to decreased weights of 0.67, 0.75 and 0.8 for each additional language respectively. 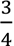 provided the most accurate approximation based on our simulated language profiles (see Figure.1) and thus was the weighing function we settled on.

### 2.3 Validating our proposed measure

While these formulas were designed based on theoretical and empirical foundations from prior research, their practical applicability required experimental validation. Thus, we next validated their robustness by testing them on two large, real-world datasets from different populations.

*Dataset 1* included language background information from 131 healthy (mostly young) monolinguals and bilinguals who completed a language questionnaire derived from LEAP-Q (Marian et al., 2007) and Language Use Questionnaire (LUQ; Kastenbaum et al., 2019; Marte et al., 2025). *Dataset 2* consisted of 139 older bilingual individuals with aphasia, who completed the LUQ. We chose these datasets to ensure that our measures can capture variance across both young/typical and older/clinical populations, enhancing their generalizability.

#### 2.3.1 Is including only Age of Acquisition and Proficiency enough?

In recent years, language background questionnaires have become increasingly detailed, collecting extensive information about participants’ linguistic history, including exposure and proficiency across different age ranges, education levels, and familial language backgrounds.

While these additions provide a more granular view, we propose a simpler approach—a formula that relies on only two key variables, under the assumption that they sufficiently capture the essential variance in language background. To test this assumption, we conducted multiple Pearson’s correlation analyses across all language background factors in both datasets (excluding post-stroke factors). Even after correcting for false discovery rate across multiple comparisons, we found that all language background variables were significantly correlated with each other in both datasets, indicating that they largely captured the same underlying variance (see Figure 2).

**Figure 2.**
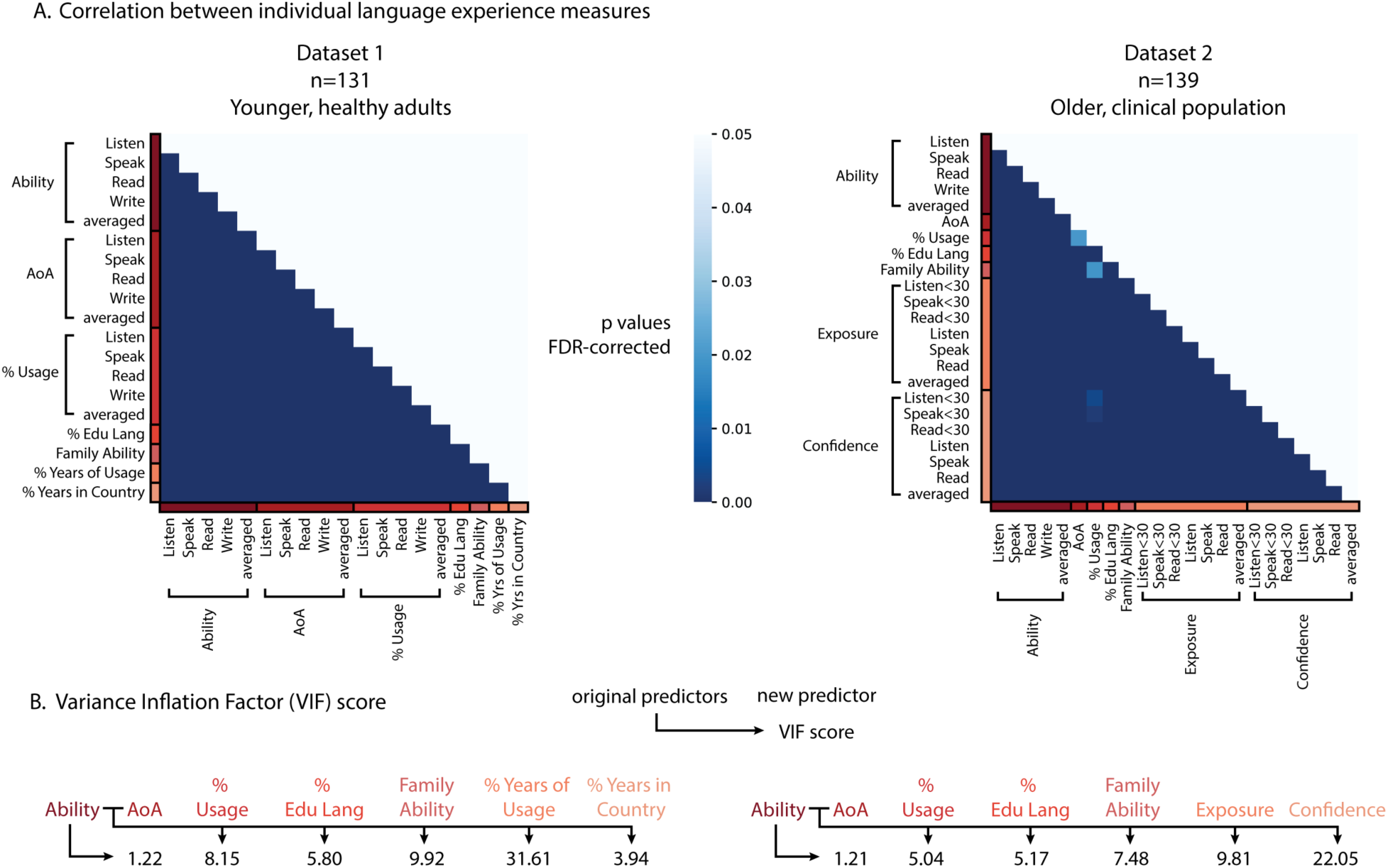
Relationship between different language background predictors. **Panel A** shows Pearson’s correlation between each pair of language background measures. All measures are highly correlated (*p*<0.05, FDR-corrected). **Panel B** shows Variance Inflection Factor (VIF) scores. AoA: age of acquisition. % Usage: percentage of time using the language, averaged across different age brackets. % Edu Lang: percentage that the language is used as the instruction language in school. Family Ability: averaged ability of the language in both parents and siblings, whenever applicable. % Years of Usage: percentage of years in life where the language was in use. % Years in Country: percentage of years in life spent in a country with the language as their official language. Exposure: averaged language exposure across different age brackets. Confidence: averaged level of confidence across different age brackets. <30 indicates that the measure is only considering the profile before age 30.

To confirm that only AoA and Ability contribute unexplained variance—making them the essential factors for our dominance and multilingualism formulas—we calculated the Variance Inflation Factor (VIF) among the language background measures. VIF quantifies multicollinearity: if adding a new predictor to the model results in a VIF score above 5, it indicates severe redundancy, meaning the new variable contributes little unique variance beyond what an existing predictor already captures.

Here, we examined whether any additional language background variable accounted for meaningful variance beyond self-rated ability. First, we tested AoA and found that in both *Dataset 1* and *Dataset 2*, it could account for variance over and above that accounted for (*Dataset 1*: VIF = 1.22; *Dataset 2*: VIF = 1.21). Subsequently, we tested the rest of the predictors on both self-rated ability and AoA. Across both datasets, the only predcitor that had a score below 5 is the percentage of years spent in a country where the language of interest is the official language (% Years in Country, VIF=3.94, *Dataset 1*), which could be an indicator of language immersion. All other predictors showed substantial multicollinearity, with VIF values well above the threshold (e.g., percentage usage across life (VIF = 8.15 and 5.04, respectively), percentage of language used in Education (VIF = 5.80 and 5.17), averaged ability of the Family (VIF = 9.92 and 7.48).

These results confirm that beyond AoA, no additional predictor provided independent explanatory power (except for % of Years in Country, but see Validation and Discussion below). Therefore, our decision to use only self-rated ability and AoA is empirically justified, as adding more variables would create redundancy without contributing meaningful new information.

#### 2.3.2 Validation against multidimensional (PCA) approaches

Finally, we compared the performance of our language dominance measure against multidimensional approaches that have emerged in recent years (Chen et al., submitted; Kastenbaum et al., 2019; Marte et al., 2025). Our measure offers a simplified and streamlined approach by deriving the dominance score from the two key dimensions that account for the most variance across language background questionnaires. However, an alternative approach could be to leverage the high-dimensional language background data and apply dimensionality reduction to derive dominance measures through Principal Component Analysis (PCA).

Here, we compared our language dominance measure to a PCA-based dominance calculation in *Dataset 1* (n=102 excluding monolinguals) and *Dataset 2*. Following prior work, we calculated the difference between English and participants’ L2 across all variables in each language background questionnaire (LEAP-Q and LUQ). We standardized the data to have a mean of 0 and a standard deviation of 1, ensuring comparability across all dimensions. Then, we derived a PCA-based dominance score by projecting individual data points onto the first principal component. To interpret the signage and magnitude of the projected values, we scaled them linearly between −1 and 1 based on the projection value of two simulated monolingual profiles on both sides. Like in our language dominance measure, −1 and 1 in the scaled PCA represents monolinguals on both sides, and 0 indicates a perfectly balanced bilingual.

We used two approaches to assess the similarity between the two language dominance measures. First, we calculated Spearman rank correlations between the PCA-based dominance score and our proposed dominance measure. Second, we categorized participants as either English-dominant or L2-dominant based on both measures: scores between −1 and 0 indicated L2 dominance, while scores between 0 and 1 indicated English dominance. A strong correlation and a low percentage of classification differences would suggest that our simpler, theoretically grounded measure closely aligns with the more complex, PCA-based approach.

In both datasets, the two dominance measures were strongly correlated (*Dataset 1: r* =.96, *p* < 0.001; *Dataset 2: r* =.94, *p* < 0.001; Fig 3), demonstrating that collecting large amounts of language background data and applying multidimensional analyses does not yield a more accurate dominance calculation. For the categorical classification, 95.1% of the participants in *Dataset 1* and 94.2% of the participants in *Dataset 2* were consistently classified by the two dominance measures. Further inspection of the people who were classified differently showed that they were very close to a perfectly balanced bilingual (value of <0.03 in at least one of the two measures), making the classification into categorical dominance groups essentially moot.

**Figure 3.**
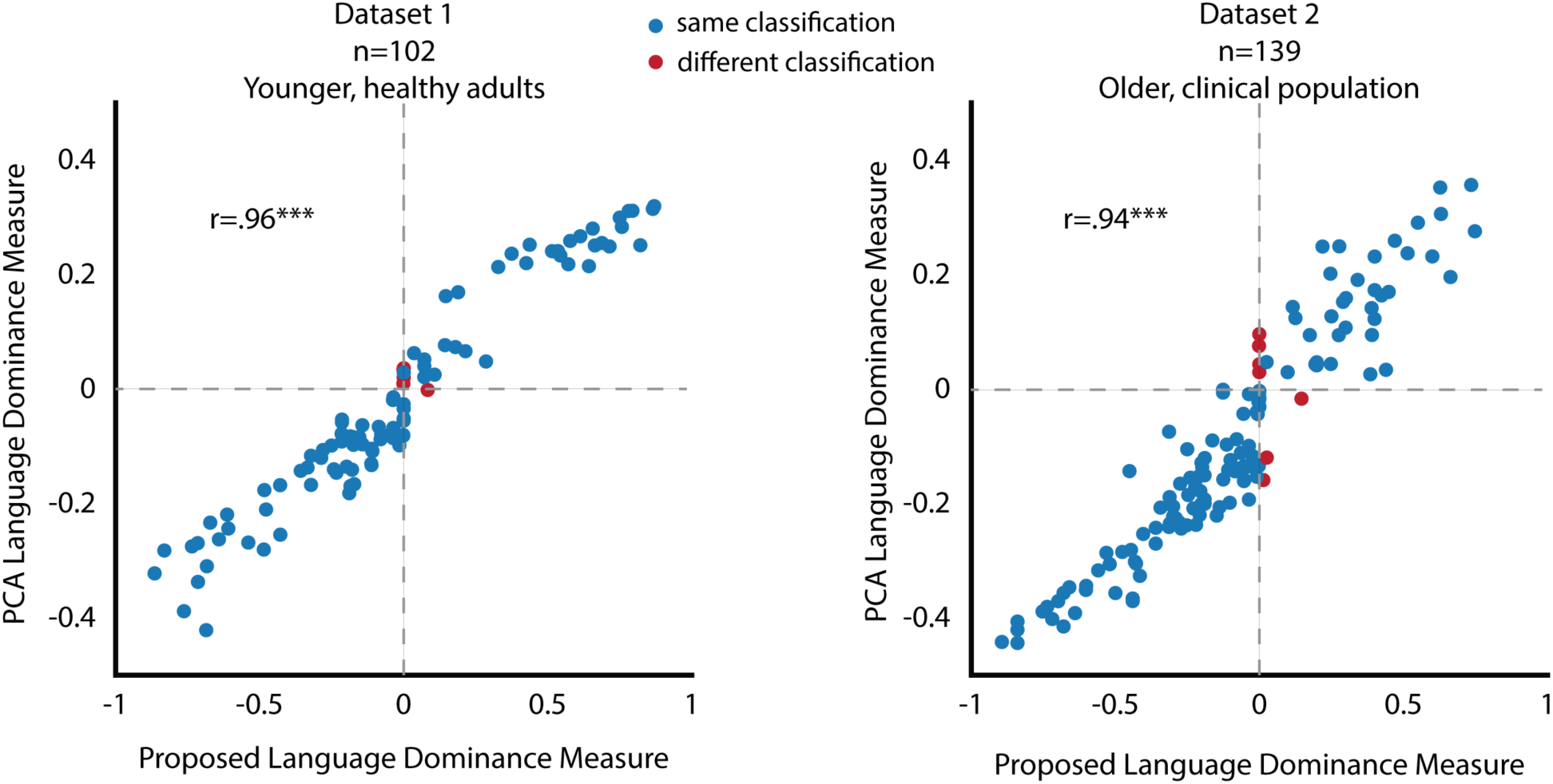
Comparison between our proposed theoretically grounded and PCA-based language dominance measures. blue dots represent data points classified in the same dominance group across both measures, while red dots indicate data points assigned to different dominance groups in the two measures. The two language dominance measures produce nearly identical results, showing a near-perfect correlation with only a few data points classified differently.

## Discussion

In this study, we addressed a long-standing challenge in bilingualism research: the accurate and standardized assessment of language dominance and degree of multilingualism. While existing assessment tools provide valuable insights into linguistic background, they often rely on heterogeneous methodologies that limit cross-study comparability. Our proposed formulas offer a computationally streamlined approach to measuring language dominance on a continuous scale, providing a replicable and efficient alternative to existing methods.

One key innovation of our approach is the reliance on self-rated proficiency and AoA as the primary variables for dominance calculation. Through multiple empirical validations, we demonstrated that these two variables capture the vast majority of variance in bilingual language profiles, rendering additional background measures largely redundant. This supports the decision to prioritize a parsimonious model that reduces participant burden and enhances methodological consistency without sacrificing predictive accuracy. One could further refine the measure to incorporate the percentage of years in life spent in a country with the language as their official language, which is the only other measure that showed a VIF score lower than 5 and suggested low multicollinearity. This variable can be a measure of language immersion that is modeled for language attainment in Hartshorne et al. (2018). However, given the high similarity between the language dominance derived from our measure and the PCA approach where the information was available, we do not think it is necessary for the current purpose.

Furthermore, our logistic transformation of AoA introduces a biologically informed adjustment that accounts for the diminishing impact of later language exposure on ultimate proficiency. This function, derived from a large-scale study of bilingual language attainment (e.g., Hartshorne et al., 2018), allows us to model a realistic trajectory of proficiency development, ensuring that the dominance measure reflects both self-reported ability and age-related constraints on language learning.

The strong correlation between our dominance measure and PCA-based dominance scores (r = .94 and r = .96, *p* < .001) demonstrates that our simple formula achieves results comparable to high-dimensional statistical methods. These findings suggest that complex, multidimensional statistical techniques do not necessarily offer superior assessments of language dominance; instead, a well-calibrated, theoretically motivated formula can yield equally reliable results while maintaining ease of application.

Beyond dominance calculations, we also introduced a multilingualism score that situates bilingualism/multilingualism within a broader, continuous spectrum of language experience. By weighting each language’s contribution to an individual’s linguistic repertoire, our approach acknowledges the differential impact of adding additional languages to one’s language profile. This measure aligns with recent shifts in the field toward viewing bilingualism as a gradient rather than a categorical variable, providing a quantitative framework for researchers investigating multilingual populations.

Overall, we propose a standardized and replicable method for measuring language dominance and multilingualism, grounded in both empirical evidence and biological constraints. By offering a computationally efficient and easily implementable framework, that does not rely on labor intensive language background collecting practices, our approach has the potential to enhance cross-study comparability and methodological consistency in bilingualism research, providing a valuable tool for future investigations into multilingual language experience.

## Acknowledgements

This research was supported by NIH grant R00DC019973 and NSF grant 2446452 to EBE.

